# Whale shark residency and small-scale movements around oil and gas platforms in Qatar

**DOI:** 10.1101/2023.11.15.567314

**Authors:** Steffen S. Bach, David P. Robinson, Mohammed Y. Jaidah, Simon J. Pierce, Prasad Thoppil, Christoph A. Rohner

## Abstract

Artificial structures in the ocean can influence the movements and residency of migratory fishes. Whale sharks seasonally aggregate near oil and gas platforms in Qatar to feed on fish spawn, creating one of the largest aggregations for the species. We used passive acoustic telemetry to examine their fine-scale movements, residency, and seasonality and investigate whether the platforms influence their habitat use in the area. Tags had a mean retention of 161 ±186 SD days and 32 of the 117 tags were recorded in multiple seasons in the acoustic array (21 stations). Most detections were recorded during the season that was established with other methods from May to September, confirming that this whale shark aggregation is seasonal. Whale sharks stayed up to 77 consecutive days in the array (mean = 16 ± 12.51 days) and had a mean residency index R_max_ of 0.31, highlighting the importance of this site to their ecology. While most detections were made at a receiver near a platform, other platforms had few detections and the distance from the centre of the aggregation was the main explanatory variable in a GLM, indicating that the platforms do not influence the whale shark’s habitat use. Instead, they moved with the current during the morning, when they feed on fish eggs at the surface which also float with the current, and swam against the current in the late afternoon and at night to be at the presumed fish spawning site again in the early morning. Our results highlight the importance of this small feeding area for whale sharks which face a high threat level in the region.

## Introduction

Marine pelagic species live in a vast environment that, despite seeming homogenous, is characterised by spatiotemporal variation in biological, chemical, and physical parameters [1]. These heterogeneous conditions result in a patchy resource landscape, with biological diversity and productivity concentrated in relatively small areas, such as in fronts, near seamounts and oceanic islands, or even around anthropogenic structures [2–4]. These productive areas influence the distribution of pelagic animals and can be preferred aggregation sites [5]. These aggregation sites are commonly important to the ecology and social behaviours of marine species [6–9]. Such aggregations may, however, also increase a species vulnerability to human-induced threats [10], and targeted fishing of aggregations can result in severe population declines [11,12]. Fish aggregating devices (FADs) exploit the same principle and target commercially important species, such as tuna [13]. Managing the exploitation of aggregation sites, and ensuring their persistence, is therefore an important consideration in the conservation of threatened pelagic species.

Whale sharks *Rhincodon typus* aggregate in ∼30 currently known hotspots, termed ‘constellations’, in tropical and subtropical waters around the world [14,15]. Aggregation sites are important to whale sharks that are otherwise solitary and wide-ranging, and these sites are thus often focal areas for conservation management of this Endangered species. Many constellations are seasonal and are driven by high prey availability [16,17]. Feeding in high-density, but ephemeral, prey patches is likely to be crucial in whale shark nutrition, as they may undergo long periods of fasting between productive feeding bouts [18]. Examining their residency and small-scale movements within an aggregation site can thus improve our understanding of the importance of these sites and aid in their protection.

Although whale sharks are often seasonally sighted at the surface in constellations, passive acoustic telemetry has revealed that some whale sharks may be present at other times, emphasising the importance of sightings-independent methods to examine seasonality and habitat use. The percentage of sharks that remained in the constellation areas throughout the year varied among different study sites, highlighting the need for site-specific investigations [19–21]. Satellite-tagged whale sharks may move 1000’s of kilometres away from aggregation sites [22], though many individuals return to constellations in subsequent seasons [15]. Their fine-scale movements and temporal shifts in habitat use within constellations are less understood. Off Mafia Island in Tanzania, the whale shark core habitat shifted over the year, from feeding areas close to shore during the main sighting season, to a few kilometres offshore between seasons [23]. Elsewhere, studies have typically focused on seasonality and residency rather than fine-scale movements.

Segregation by sex and size is a ubiquitous characteristic among whale shark constellations. There is a substantial bias towards male sharks at most locations, and small (0.6–3 m total length TL) or large (>8 m TL) individuals are rarely seen at these sites [14]. One exception to the sexual segregation has been documented in the Red Sea, where juvenile females and males occur in an even ratio. Passive acoustic telemetry and satellite tagging showed no differences in residency or wider-scale movements between the juveniles of both sexes [20]. Another exception is the offshore island of St Helena in the mid-Atlantic, where mostly large, mature whale sharks are sighted [24,25]. At present, it is unclear whether these large sharks display temporal changes to their habitat use within constellations, or how their small-scale movements compare to those of smaller individuals.

One of the largest whale shark constellations in the world has been documented in oil and gas fields off the coast of Qatar in the Arabian Gulf. Whale sharks seasonally gather here from May–Sep and primarily feed on freshly-spawned fish eggs from mackerel tuna (*Euthynnus affinis*) [26]. Densities of up to 100 sharks in a 1 km^2^ area have been reported [26]. More males (69%) than females are seen off Qatar, and there is an unusually large proportion of mature males (63% of males) and even some large females, unlike the juvenile bias in other regional hotspots [15], making it possible to examine small-scale habitat use and residency across a broad size range of sharks in a population for the first time.

The whale shark constellation in Qatar occurs in an area of active oil and gas extraction known as the Al Shaheen oil field and the North field. The first reports of this constellation were made by workers on oil and gas platforms, and it is unknown whether whale sharks aggregated in this area before the establishment of platforms [26]. Platforms elsewhere have also reported the presence of whale sharks [27,28] and more generally attract migratory animals, such as tuna, billfish, and sharks [2]. To test whether it is the platforms specifically that attract the sharks, or whether the platforms are simply in the general area in which fish eggs are available, observations at and around several platforms are needed.

To that end, we used passive acoustic telemetry to examine whale shark movements within their aggregation site in Qatar. Receivers were placed near platforms and in open water, to test the association between whale sharks and these structures. We also investigated whale shark residency and the seasonality of this constellation, and examined the core habitat and the timing of their movements between stations in the receiver array.

## Methods

### Study area

The Arabian Gulf, ‘the Gulf’ hereafter, is an almost enclosed sea that is connected to the Gulf of Oman and the Indian Ocean by the narrow Strait of Hormuz in the east (Fig. 1A). Located in the arid sub-tropics and surrounded by deserts, the Gulf is one of the warmest and most saline waters on earth [29,30]. Surface water temperatures fluctuate strongly across seasons, from 20°C in winter to up to 35 °C in summer in the deeper, central areas, where whale sharks have been sighted around oil and gas platforms [26,31]. Platforms in the study area are located ∼80 km north-east off the coast of Qatar, where water depth ranges from 50–70 m (Fig. 1A). The first fixed platforms related to the Al Shaheen oil field were established in 1996, while the latest was put in place in 2009. There are 34 structures in total, with some interconnected through gangways, resulting in 9 separate platform locations. The subsurface section of the platform structures occupy on average an area of 2,500 m^2^. In this area, whale sharks have mostly been sighted between May–September, here termed the “season” [26].

**Figure 1.**
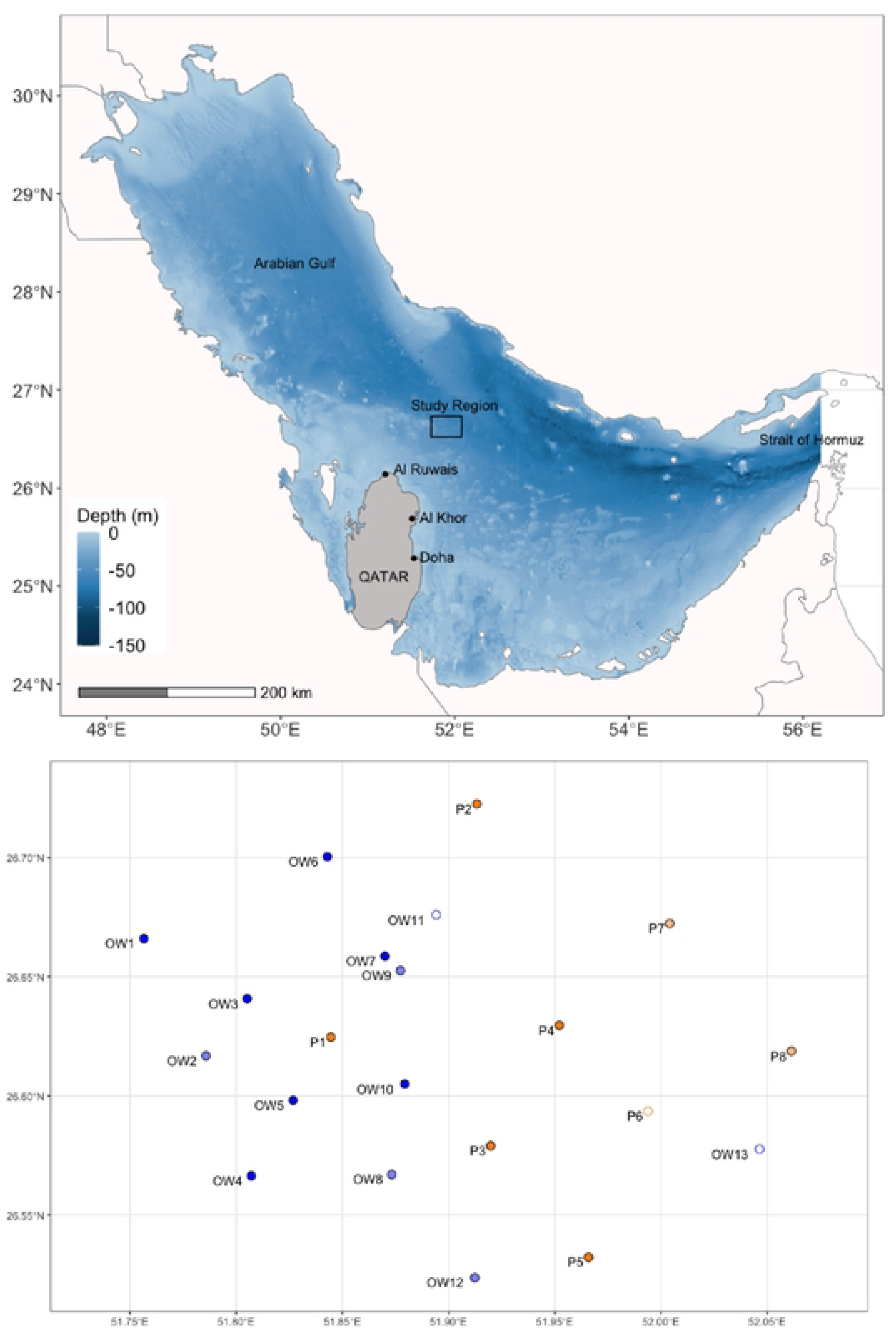
A) Regional overview map. B) The acoustic receiver array off Qatar with platform­ associated stations (Pl-PS, orange) and open water stations (OW1-0W13, blue). Empty circles indicate stations that were lost before the first data download, and the light colours represent stations that were only used in the first season (2012/13).

### Acoustic array

Acoustic monitoring equipment was deployed and whale sharks were tagged during 49 boat-based surveys between 2012–2016. A 10-metre vessel with twin 250 hp engines was used for surveys. The point of departure was the port of Al Khor or Al Ruwais on the north coast of Qatar (Fig. 1). Permissions for fieldwork and data collection on whale sharks were given by the Qatar Ministry of Municipality and Environment. Analyses were conducted in R version 4.1.2 [32].

Acoustic receivers (Vemco, VR2W) were deployed at 21 stations (Fig. 1B). To test whether whale sharks had an affinity for underwater structures, 8 receivers were deployed within 500 m of platforms (named P1–P8, east to west) and 13 receivers were located away from platforms in open water (OW1–OW13). In the first season (2012), the array comprised 12 stations at 7 platforms and 5 open water locations. The array was then adjusted from the second season onwards (2013–2016) to remove stations with few detections and add stations around P1 that recorded the most detections (74%) in the first season. There were 12 stations at 5 platforms and 7 open water locations from 2013 onwards.

Stations were in deep water (>50 m) and required acoustic releases to retrieve receivers. We attached receivers to a cement anchor (>20 kg) with an acoustic release (SubSeaSoncic AR-60-E Acoustic Release Unit) and two trawl floats with a total buoyancy of 18.8 kg (600 metres working depth and 30 kg m^-1^ impact strength). To retrieve receivers, the acoustic releases were activated with a transducer (SubSeaSonic Acoustic Release Interrogator Model ARI-60), thereby detaching from the anchor and floating to the surface by the attached floats. Receivers were regularly downloaded and redeployed, but some gaps in the coverage exist due to operational constraints (Supplementary Figure 1). We focused receiver deployments on the tuna spawning season from May–Sep, when whale sharks formed aggregations [26], but also had some receivers active in other months to test if whale sharks were absent in the non-spawning season. Receivers at three stations (P6, OW11, and OW13) were lost before any data could be downloaded and were excluded from the analyses.

### Whale shark tagging

Whale sharks were encountered and tagged with acoustic tags (Vemco V16) at the surface during their feeding aggregations. We were unable to photo-ID some (23 of 125) of the tagged individuals, and it is possible that some individual whale sharks were tagged multiple times over the study if they lost their initial tag. We visually determined the sex and estimated the total length (TL) of most (82%) tagged sharks. Tags were connected with a 10–15 cm long dyneema tether to a titanium dart, which was inserted into the skin below the 1^st^ dorsal fin of the shark. In 2012 and 2013 we used a pole spear, and in 2014–2016 we used a pneumatic spear gun to deploy the tags. Tags had an expected battery life of 2,516 days but often fell out of the shark prematurely (Table 1). Tag #72 fell off the shark next to station P1 on 18 Aug 2015 so we removed all detections thereafter before analysing the data. Tag retention was defined as the number of days between tag deployment and its last detection in the array. Therefore, retention is a minimum estimate, as whale sharks likely migrated away from the array with their tags attached and did not return, or lost their tags elsewhere before returning.

**Table 1.**
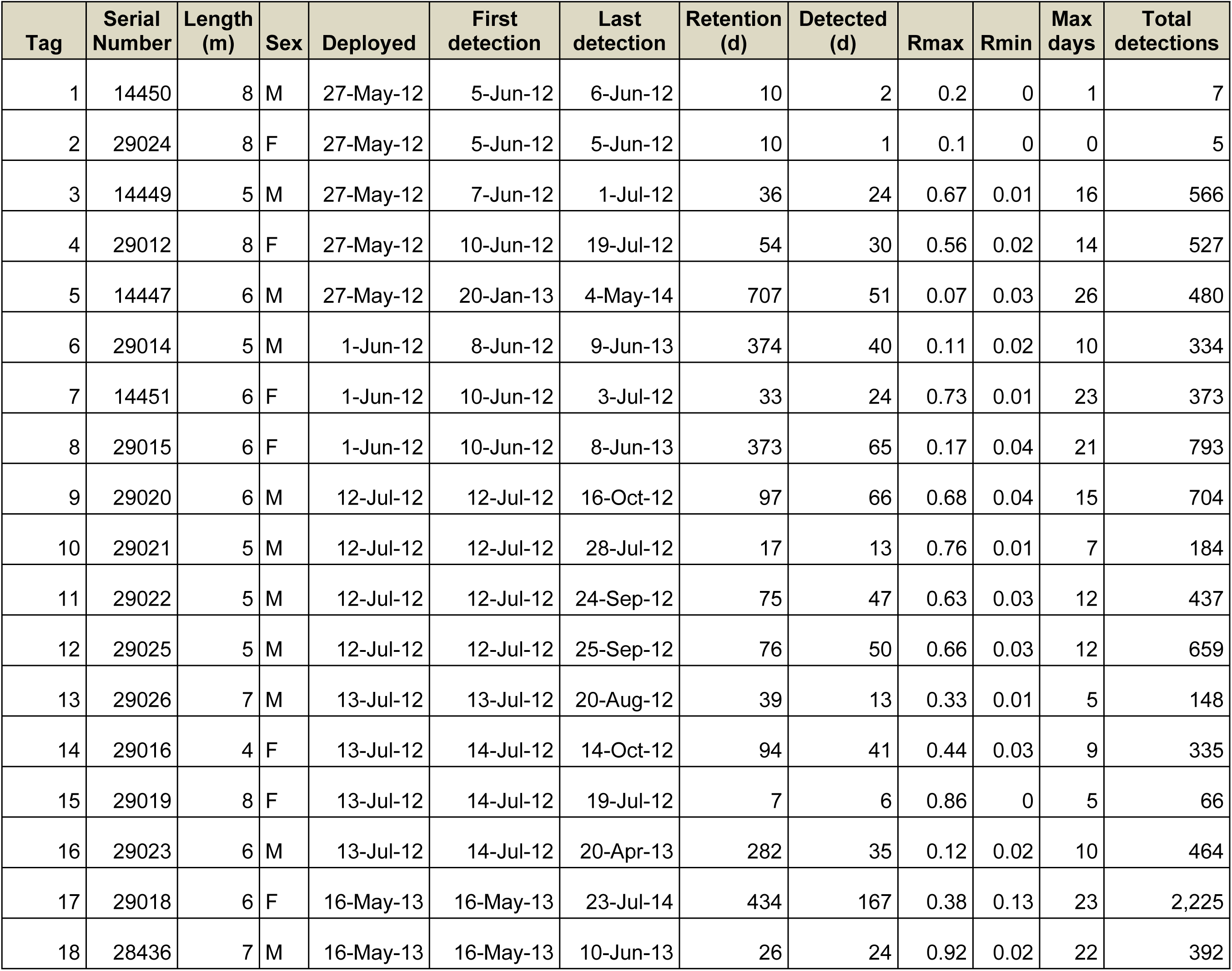
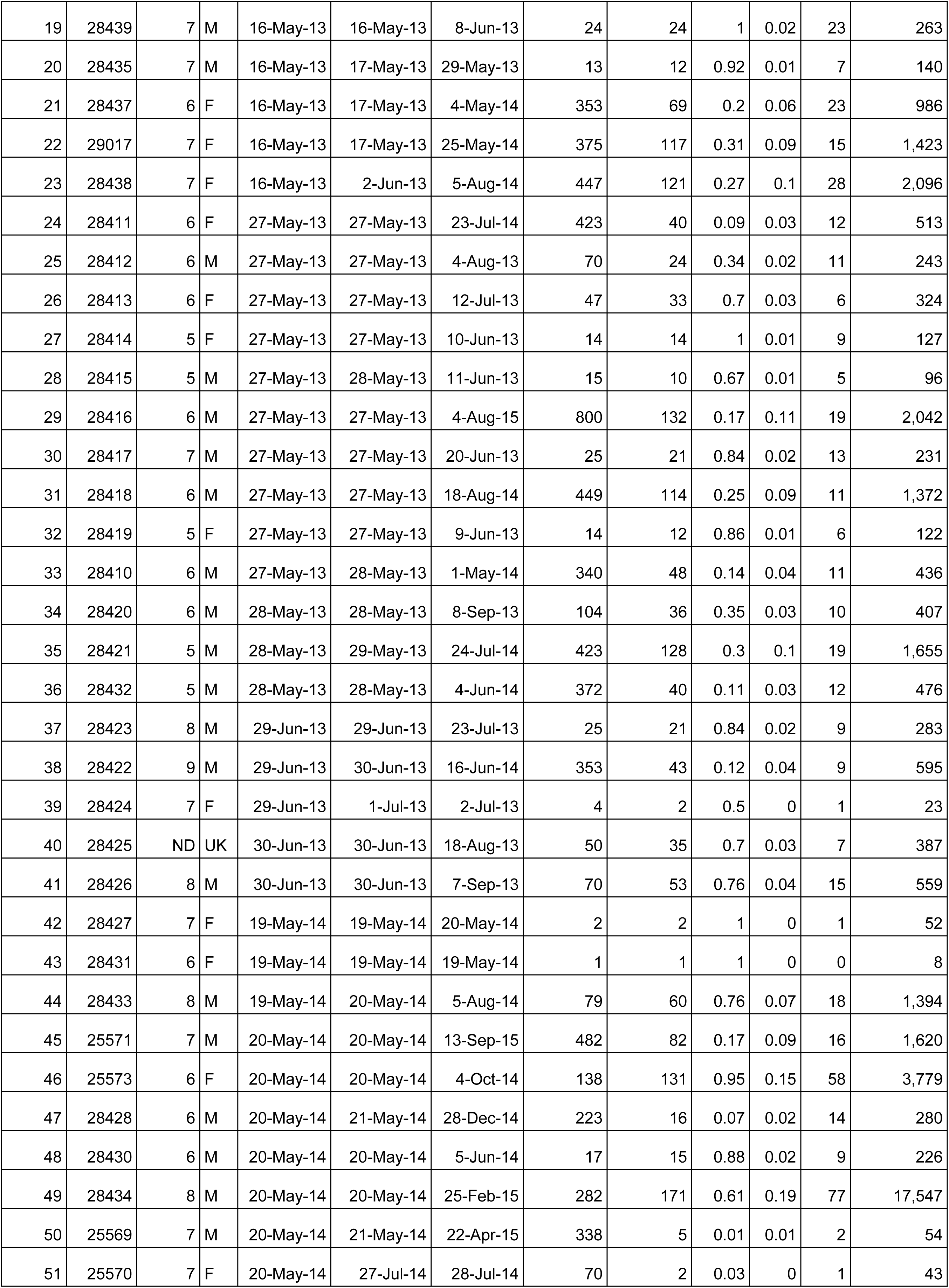
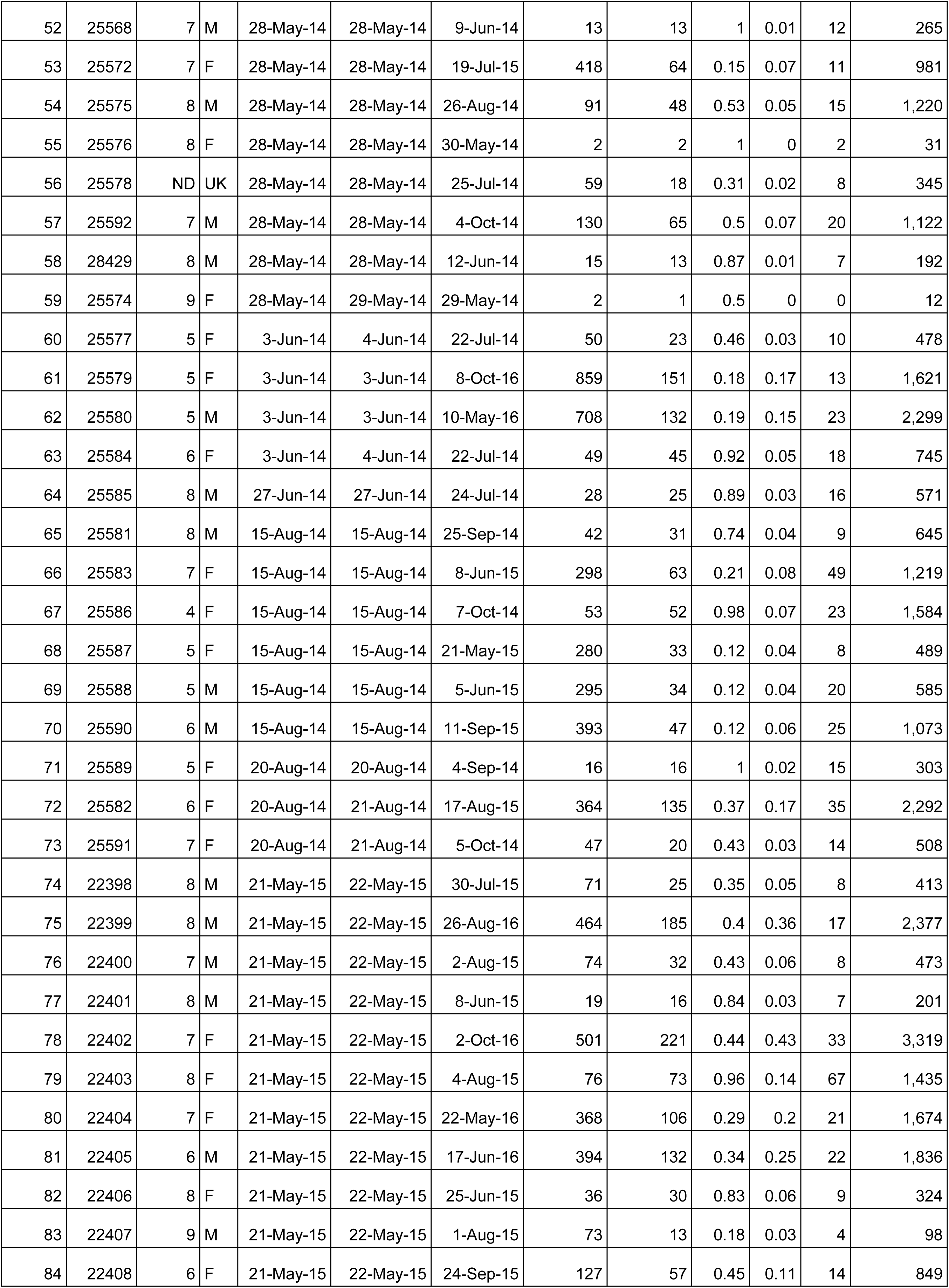
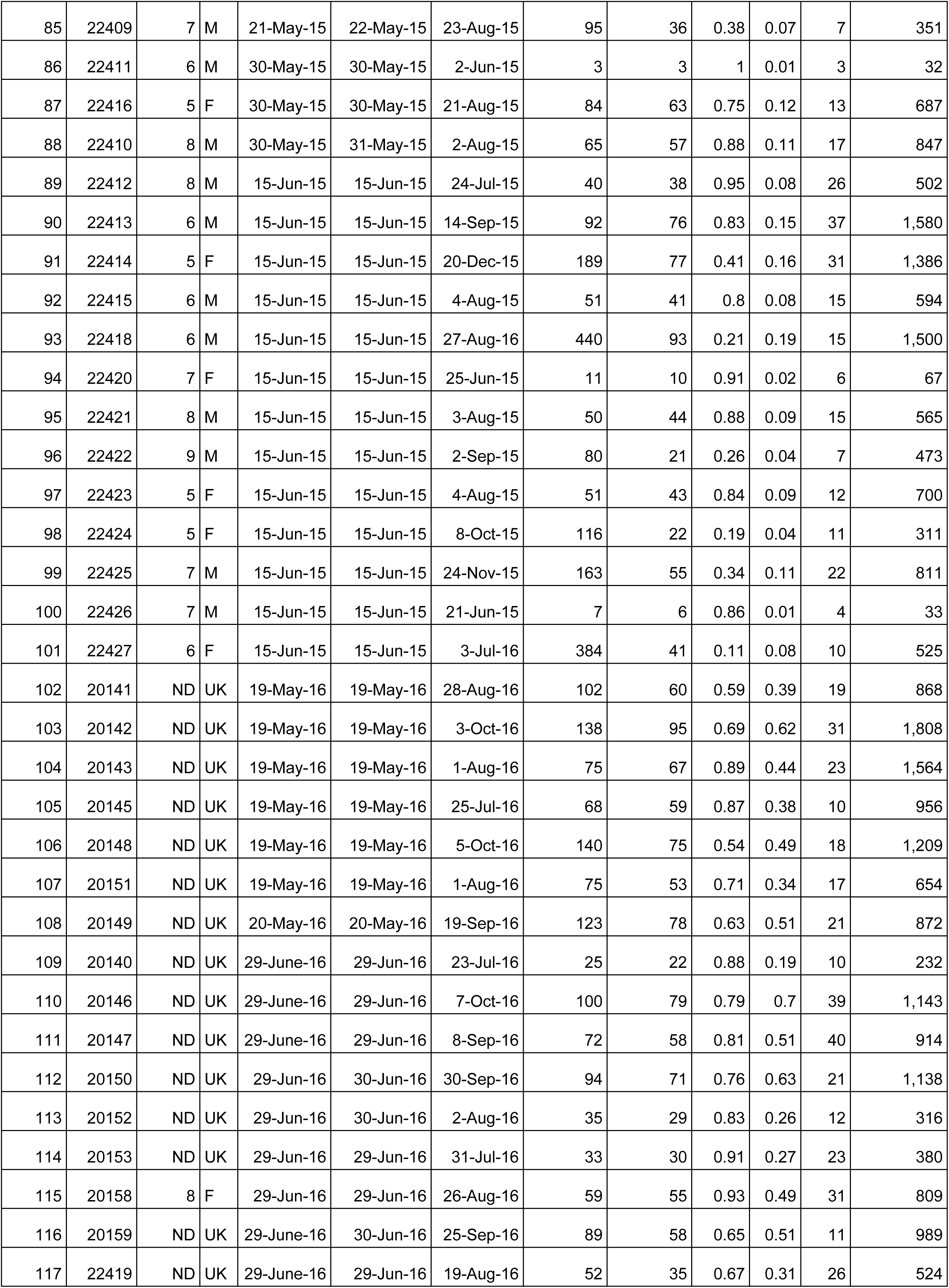
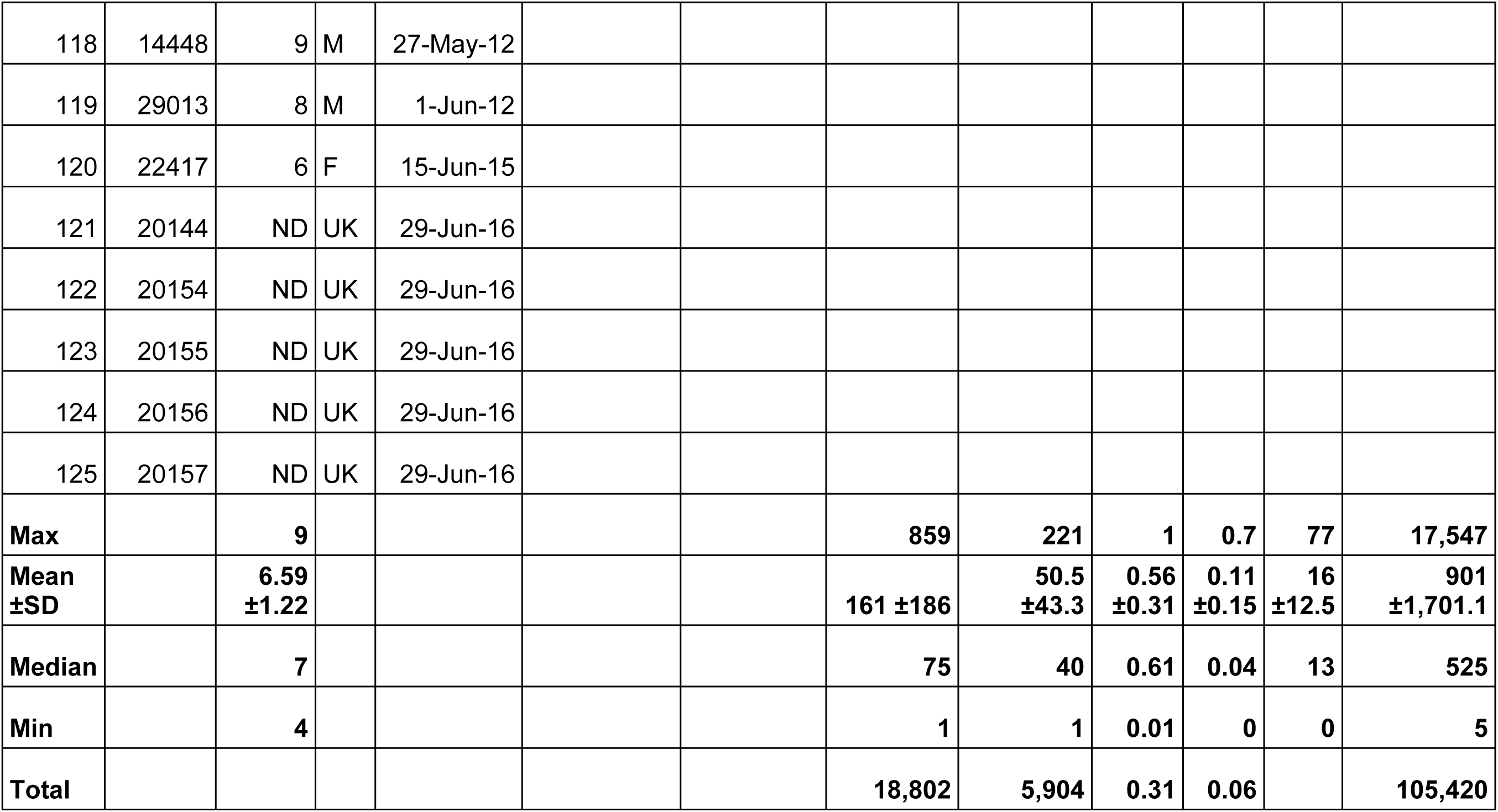
Details of all whale sharks tagged during this study. The end of the tag period for R_max_ was the date of last detection and for R_min_ it was 19 Oct 2016 when the study ended. Max days refers to the maximum number of consecutive days a shark was detected in the array.

### Residency

To investigate the importance of the feeding aggregation area to tagged whale sharks, we calculated two residency indices [20,23]: R_max_ was the proportion of days a shark was detected in the array over the number of days between tag deployment and its last detection. R_min_ was the proportion of days a shark was detected over the number of days between tag deployment and the end of the study, 19 Oct 2016, when the last receiver was recovered. Both indices were calculated for each shark individually and for all sharks combined based on the sum of detection and deployment days. We also calculated the number of consecutive days sharks were detected in the array. The number of seasons in which each shark was detected was a longer-term measure of site fidelity, although it was biased by the relatively short tag retention.

### Habitat use

The fine-scale habitat use of whale sharks within aggregation sites is poorly understood, since satellite telemetry often has a relatively large location error, and visual observations are limited by survey effort and visibility. Acoustic telemetry allowed us to examine which areas within the aggregation site were important to whale sharks, and whether their activity hotspots were correlated to platform locations. We first investigated the overall habitat use for all whale sharks combined. We used centres of activity (COA), calculated as one location per 24 h, using the *V-Track* package [33]. These COA are position estimates based on weighted means of the number of detections at stations over a time period. They can thus be spread across the array, estimating the location of animal activity at a continuous spatial scale, rather than being restricted to receiver locations only [34]. We then calculated their core (50%; “core habitat”) and extent (95%; “extent habitat”) kernel utilisation distributions (KUD) based on the COA locations using the *adehabitat* package [35].

### Association with platforms

To test whether platforms played a role in where whale sharks aggregate we constructed a generalised linear model (GLM) with the number of detections per day of recording for each station as the response variable, and the type (platform or open water) and distance from P1, being the centre of the aggregation (Fig. 1B), as predictors. We then plotted the number of detections per day of recording and distance from P1 and fitted regressions to the data. A polynomial regression had the best fit and explained 92% of the variance.

### Seasonality

We examined seasonal trends in whale shark visitation to compare with visual and satellite telemetry results that have shown that whale sharks frequent Al Shaheen between May– Sep [31,36]. We binned the total number of detections in the whole array per month and compared the May–Sep mean to the Oct–Apr mean. We also binned the number of unique tags detected per month and the percentage of detected tags from the number of available tags in that month, with the end of tag attachment taken as the last detection, as per R_max_. To investigate fine-scale variations in habitat use over the season, we calculated habitat use metrics, including the 50% and 95% core KUD, for the two seasons, and also for each month within the whale shark season from May–Sep.

### Temporal patterns

Day and night were assigned for each detection by deriving the sunset and sunrise time for each day with the *StreamMetabolism* package [37]. We wanted to assess how whale sharks move within the aggregation site over the course of the day. Field observations suggested that whale sharks drift with the current while feeding on fish eggs, which also drift, with the predominant current being from northwest to southeast. We therefore designed two transects of acoustic receivers from north-west to south-east and from north-east to south-west. We then binned detections every 30 min and plotted the relative activity of each station in the transect to investigate if the peak of activity moved with the current. The proportion of detections, rather than the number of detections, allowed us to compare stations with different levels of activity. We also derived the movement segments of whale sharks among stations, and defined a move as an occurrence of an individual being detected at a subsequent station within 3h of its first detection at the original station. We then selected the first detection at each station within a move to assess at what time of day these moves occurred.

### Currents

To assess the likely path of floating fish eggs after a spawning event, we mapped surface currents in the area based on a regional, high-resolution ocean model for the Gulf region for the months of May–October during 2012–2016, overlapping with the seasonal whale shark sightings in this study. The details of the model configuration are detailed in Thoppil and Hogan [38]. Briefly, the numerical model is the HYbrid Coordinate Ocean Model (HYCOM) with 1-km horizontal resolution and 16 hybrid layers in the vertical axis. The model domain extends northward from 22.7°N and westward from 59.4°E and has 1217 × 945 × 16 grid points. The eastern boundary is treated closed but relaxed towards the Generalized Digital Environmental Model version 3 (GDEM3) seasonally varying temperature and salinity climatology. The surface forcing for the model comes from the three-hourly, 0.5° atmospheric fields of winds, air temperature, humidity, precipitation, and solar radiation. Surface salinity is being relaxed to the GDEM3 climatology. This configuration of the Gulf model (no tidal forcing) has previously been used for other studies [38–40]. These studies have demonstrated the model’s ability to reproduce many salient features of the circulation and water masses in the Gulf, including the evolution of a series of cyclonic eddies during summer.

## Results

A total of 125 acoustic tags were deployed on 44 females, 59 males, and 22 sharks of unknown sex (Table 1). Tagged whale sharks ranged from 4–9 m in total length, with a mean of 6.6 m (± 1.22 m SD). Males (6.8 ±1.2 m) were larger than females (6.3 ±1.2 m; t = 1.995, df = 93.5, p = 0.049). Eight tags were never recorded on the receivers, and most of these tags were deployed towards the end of the study in 2016 (Table 1). Tag retention, from deployment to the last detection, ranged from 1–859 days, with a mean of 161 ±186 days and a median of 75 days. The entire acoustic array reported a total of 105,420 detections from 117 tags over the duration of the study, from 5 June 2012 to 8 October 2016 (Table 1).

### Residency

Tagged whale sharks had a high residency in the acoustic array while their tags were attached. The highest maximum number of consecutive days a shark was detected in the array was 77 days, with a mean ± sd = 16 ± 12.51 days for all sharks. Half of all sharks were detected on at least 14 consecutive days, and 12 sharks were detected on over 30 consecutive days. The residency index R_max_, the proportion of days a shark was detected in the array between tag deployment and its last detection, ranged from 0.01 to 1, with a mean of 0.56 ± 0.31 for individual sharks (Table 1). For all sharks combined, the R_max_ was 0.31. Although tags that had a short retention generally had a higher R_max_, the 32 tags detected in multiple seasons still had a relatively high combined R_max_ of 0.21. The more conservative R_min_, for which tag retention was calculated as the number of days from tag deployment to the end of the study, was much lower at 0.06 for all sharks combined (Table 1).

The residency index R_max_ varied with size, with large whale sharks (≥8 m TL; combined R_max_ = 0.51) being more resident than small (≤5 m; 0.25) and medium-sized (6–7 m; 0.26) individuals. However, large whale sharks also had a shorter mean tag retention (80.9 days) than small (191.4 days) and medium (214.6 days) individuals. Females (combined R_max_ = 0.31) had a similar residency level to males (0.27).

### Habitat use within the aggregation site

Whale sharks had a defined hotspot in their habitat around station P1 (Fig. 2). Station P1 recorded 65% of all detections within the array, followed by stations OW10 (8.5%) and P3 (8.1%) to the south-east of P1 (Table 2). When taking the variable effort into consideration, the same three stations had the highest number of detections per day of recording, although OW10 (16.4 detections day^-1^) was twice as important as P3 (8.9 detections day^-1^; Table 2). Relative numbers also showed that OW3 was more important than OW5 even though they had a similar number of total detections.

**Fig. 2.**
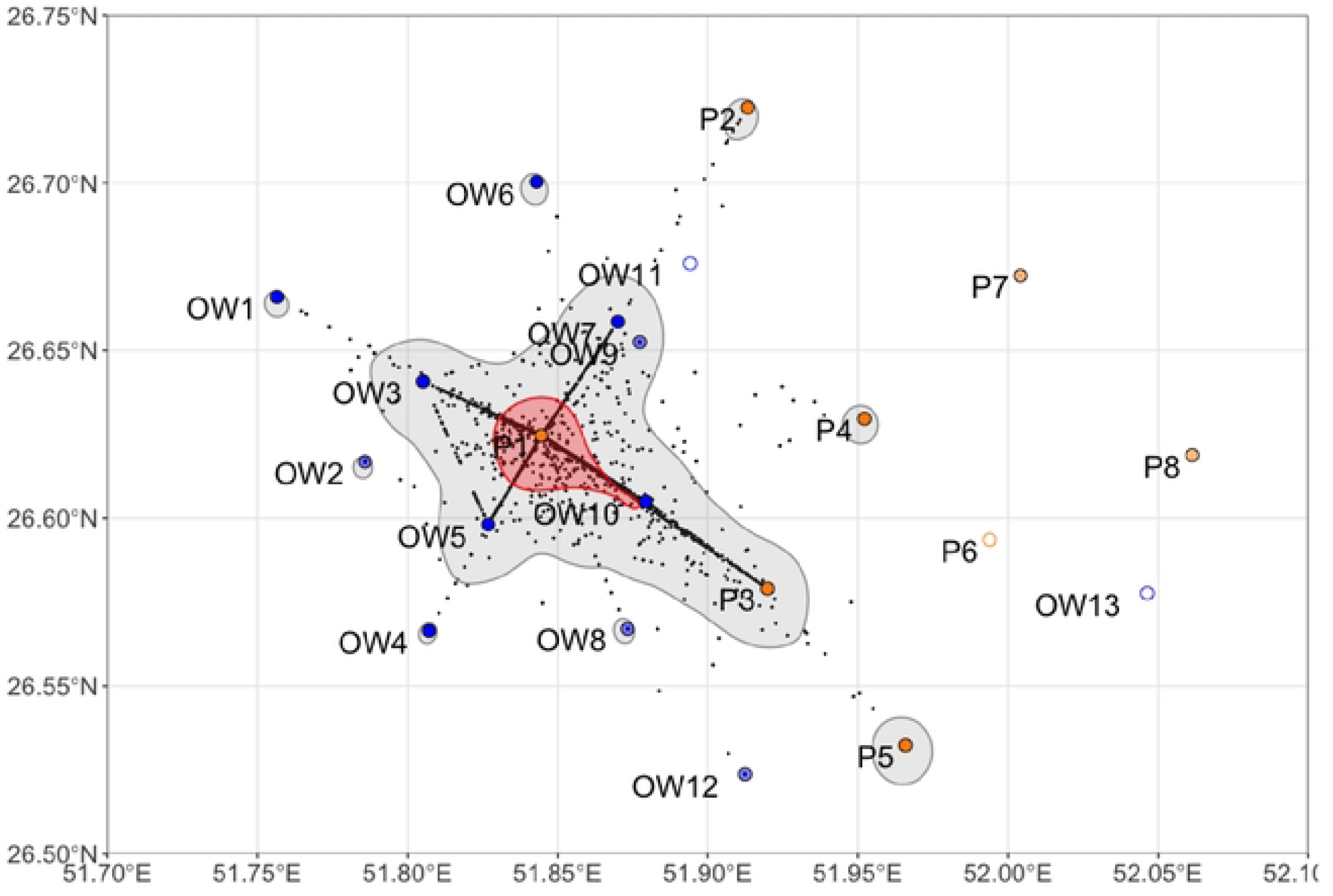
Habitat use of all whale sharks combined over the entire study period, with core (95%; red) and extent (50%; grey) kernel utilisation distributions and centres of activity per 24 h period (black dots). Stations at platforms (Pl andin open water (OW) are shown. Emptycircles indicate stations that were lost before the first data download, andthe light colours represent stations that were only used in the first season (2012/13).

**Table 2.** Activity metrics for all stations in the whale shark aggregation area.

| Station | Detections (%) | Days of recording | Days with detections | Detections per day of recording | Tags recoded | Tags per 100 days of recording |
| --- | --- | --- | --- | --- | --- | --- |
| P1 | 68,474 (65%) | 1163 | 825 | 58.9 | 114 (97.4%) | 9.8 |
| P2 | 382 (0.4%) | 951 | 18 | 0.4 | 15 (12.8%) | 1.6 |
| P3 | 8,562 (8.1%) | 959 | 357 | 8.9 | 100 (85.5%) | 10.4 |
| P4 | 757 (0.7%) | 830 | 64 | 0.9 | 45 (38.5%) | 5.4 |
| P5 | 907 (0.9%) | 961 | 109 | 0.9 | 71 (60.7%) | 7.4 |
| P7 | 2 (0%) | 318 | 1 | 0 | 1 (0.9%) | 0.3 |
| P8 | 0 (0%) | 318 | 0 | 0 | 0 (0%) | 0 |
| OW1 | 234 (0.2%) | 145 | 13 | 1.6 | 22 (18.8%) | 15.2 |
| OW2 | 86 (0.1%) | 199 | 12 | 0.4 | 11 (9.4%) | 5.5 |
| OW3 | 5,544 (5.3%) | 678 | 301 | 8.2 | 96 (82.1%) | 14.2 |
| OW4 | 163 (0.2%) | 301 | 24 | 0.5 | 22 (18.8%) | 7.3 |
| OW5 | 5,847 (5.5%) | 938 | 325 | 6.2 | 109 (93.2%) | 11.6 |
| OW6 | 365 (0.3%) | 499 | 34 | 0.7 | 29 (24.8%) | 5.8 |
| OW7 | 4,702 (4.5%) | 736 | 245 | 6.4 | 90 (76.9%) | 12.2 |
| OW8 | 266 (0.3%) | 199 | 34 | 1.3 | 12 (10.3%) | 6 |
| OW9 | 92 (0.1%) | 234 | 18 | 0.4 | 8 (6.8%) | 3.4 |
| OW10 | 8,994 (8.5%) | 548 | 369 | 16.4 | 79 (67.5%) | 14.4 |
| OW12 | 43 (0.0%) | 199 | 9 | 0.2 | 4 (3.4%) | 2 |

Most individual tags were detected at P1, where 114 of 117 tags were recorded (Table 2). Other stations with high numbers of detections (P3, OW3,5,7,10) also recorded many individual tags. The main exception was P5 at the southeast corner of the array which had few detections (0.9% of total) but recorded relatively many individual tags (71 of 117). Relative to the number of days of recording, OW1 stood out with 22 tags recorded in only 145 days.

The core habitat of whale sharks was concentrated around P1 and extended to OW10 to the south-east (Fig. 2). The extent habitat was around the core habitat, also extending to the south-east. Most stations outside of the main habitat use area also had a small, separate extent habitat surrounding them, with the exception of two platforms in the north-east and OW12 in the south.

### Association with platforms

The GLM showed that type of station (platform > open water, p = 0.002) and distance from the centre of the aggregation (p = 0.0002) influenced the number of detections per day of recording at the stations. All 7 stations at platforms combined had a mean of 14.4 detections per day of recording, while the 11 stations in open water had a mean of 5.6 detections. However, only P1 and P3 from the platform group had significant numbers of detections per day (>1.0), while 6 stations of the open water group had > 1.0 detections per day. Taken together with the higher significance level in the GLM, we pursued the distance from the centre of the aggregation and fitted a polynomial regression to the data which showed that fewer detections were made further from P1 (Fig. 3; F = 46.7, df = 13, p < 0.0001), with a plateau after ∼6 km. Stations OW10 and P3, in a south-easterly direction from P1, had more detections than the regression estimated, while nearby stations OW5 and OW9 perpendicular to that line, had fewer detections than expected (Fig. 3).

**Fig. 3.**
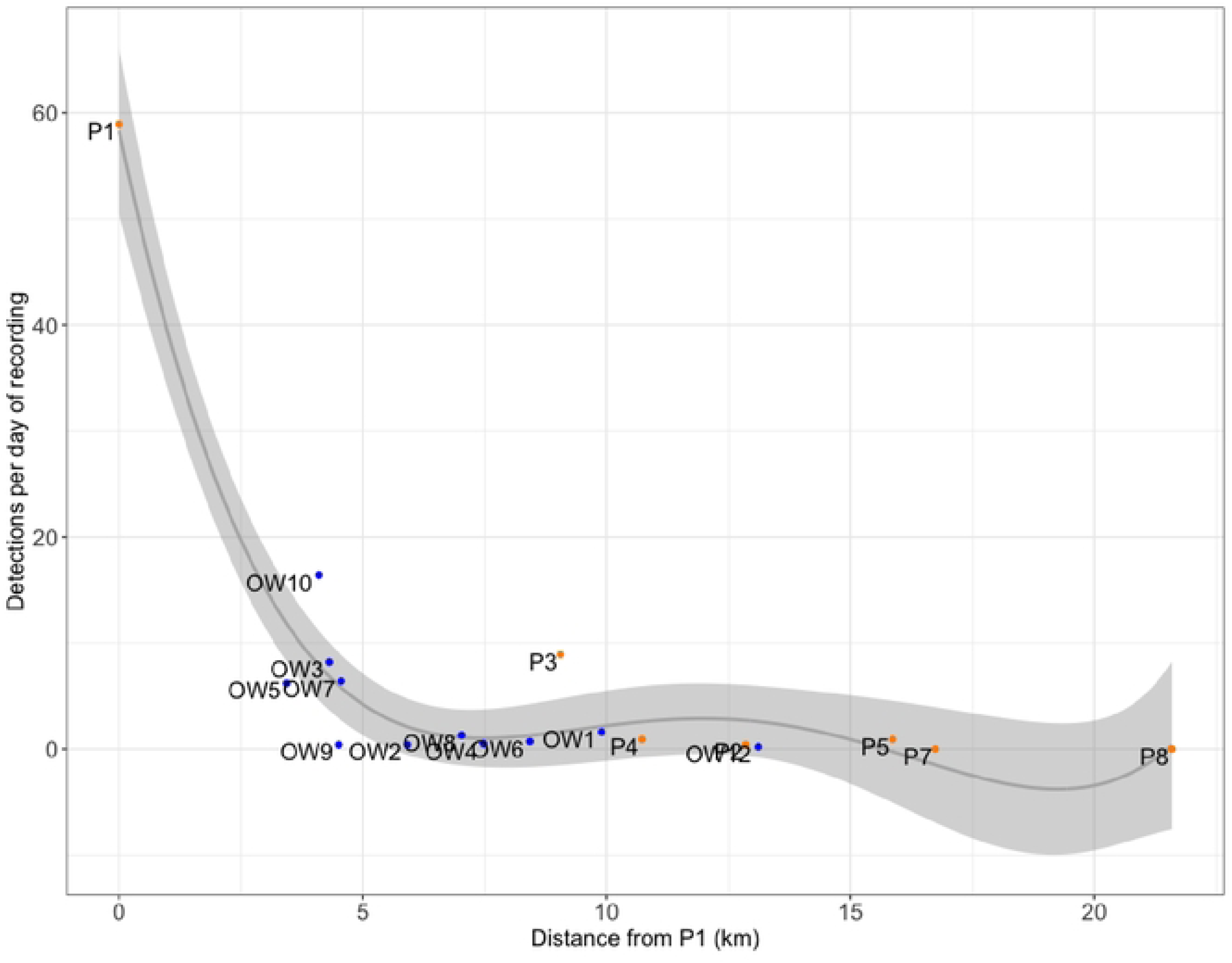
Detections per day of recording and the distance of each station to Pl, which was the centre of the activity space. Stations at platforms are shown in orange and stations in open water are in blue, with a polynomial regression line (black) and its confidence area (grey).

### Seasonality

Whale shark detections in the array were highest from May–Sep, with a peak of 28,447 detections (27% of all detections) in July (Fig. 4). Whale sharks were detected in every month of the year, however at lower levels between Oct–Apr (8.7% of detections) than between May–Sep (91.3%; t = 5.99, df = 4.46, p = 0.003). The number of individual tags detected in the array had a similar seasonal trend to the number of detections, with a maximum of 115 tags recorded in July. When taking tag loss into consideration, and calculating the percentage of tags detected from all tags available for detection as per R_max_, the seasonal trend was similar, with up to 91% of tags detected in July. One exception was in April, when a high percentage of tags was recorded (77%), but overall few tags (27) and detections (2.9%) were recorded in that month (Fig. 4). Trends were similar in each individual year from 2012–2016, with some minor variation at the beginning and end of the season. A relatively high number of detections were made in May 2013 (20.4% of that year) and May 2016 (15.3%), as well as in September 2012 (27.7%) and October 2014 (9.9%). Activity spaces varied between the main whale shark season (May–Sep) and the rest of the year (Oct–Apr), with a less defined habitat use in the latter. Within the main season there was no variation in habitat use among months. P1 was the centre of a small core habitat in all months from May–Sep (Supplementary Fig. 2).

**Fig. 4.**
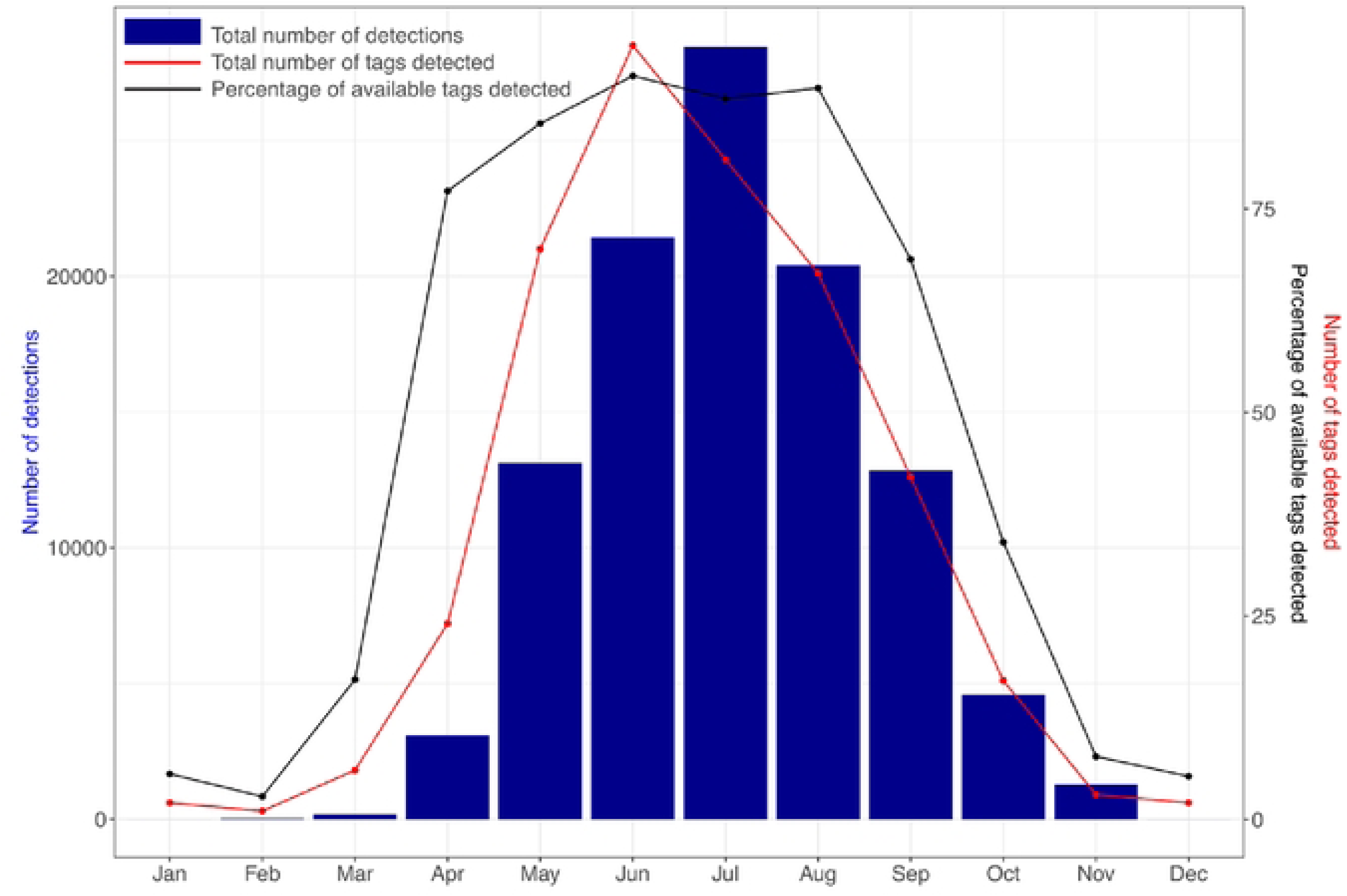
The overall number of detections (blue bars) made in the whole acoustic array by all tags combined and summed per month, with the number of tags (red) and the percentage of tags from the available tags (black) detected in the array per month.

### Temporal trends

Whale shark detections were made throughout the day (n = 53,819) and night (n = 51,601), with only a slight trend towards fewer detections from early morning to midday. There were some interesting spatial differences in the temporal trends within the array. In their core habitat most detections were recorded at P1 (37%) between midnight and 6 am, followed by a sharp decline to the minimum at 11:30 am and a slow rise to midnight (Fig. 5). Detections for five stations in a transect line from north-west to south-east (stations OW3, P1, OW10, P3 and P5) binned by 30 minutes showed that the peak of shark detections moved from north-west to south-east over the course of the day (Fig. 5).

**Fig. 5.**
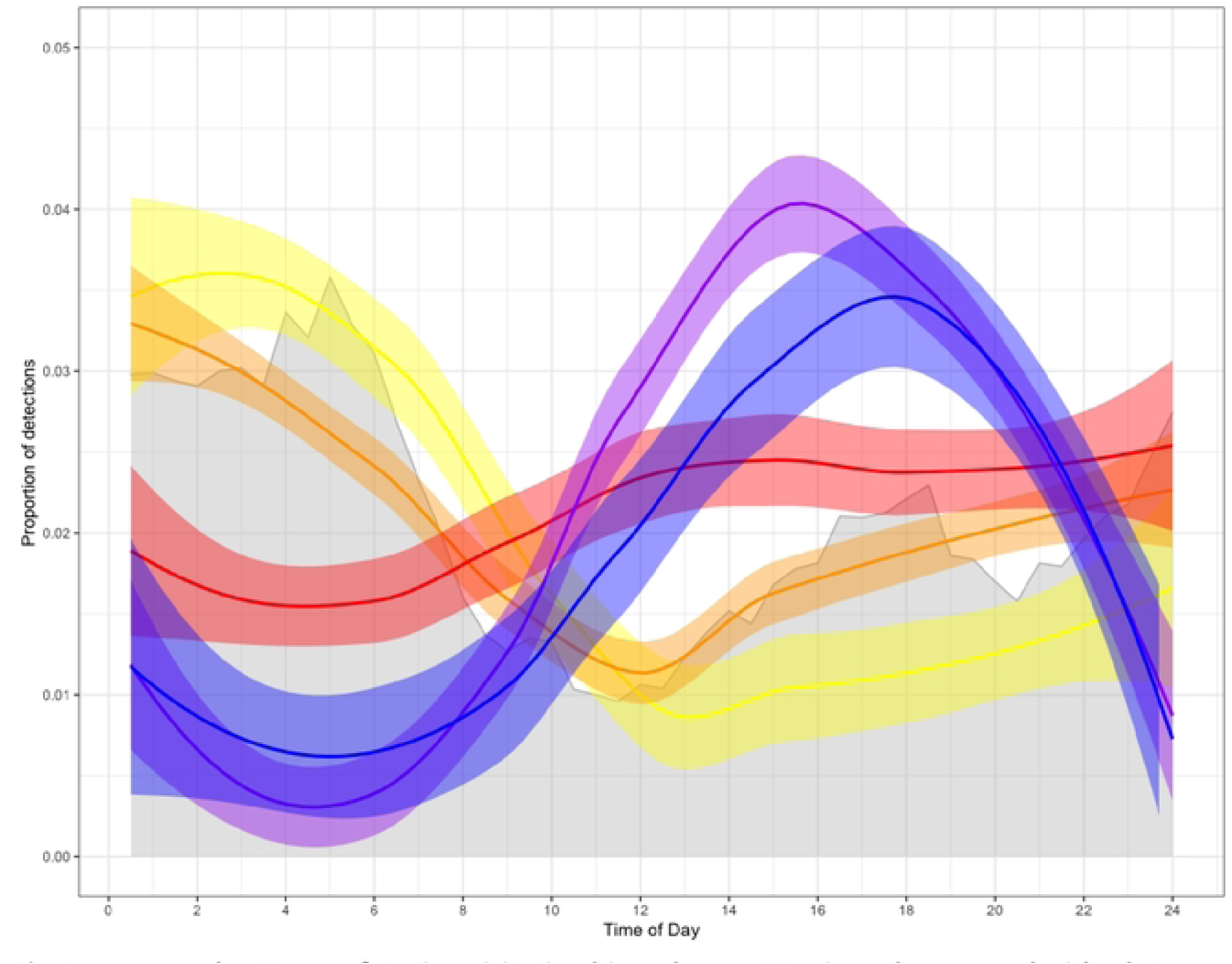
Temporal patterns of station visitation binned every 30 min and expressed with a loess smoother along a transect from north-west to south-east, with station OW3 (yellow), Pl (orange), OWlO (red), P3 (purple) and PS (blue). The raw 30-min binned data for Pl is shown in grey.

Individual moves between stations were most frequent from OW10 to P1 (n = 239), followed by moves from OW3 to P1 (n = 94), OW5 to P1 (n = 93) and P1 to OW10 (n = 86; Table 3). There was again a clear pattern with the predominant north-west to south-east current direction, with sharks mostly arriving at the downstream station during the morning (P1, OW5, OW10) and the afternoon for the stations further downstream (P3, P5; Table 3). Sharks mostly swam against the predominant current during the evening and night.

**Table 3.** The number of occurrences of moves between stations within a 3 h window (n), and the relative frequency of first detection times at the last station binned into night, morning, afternoon, and evening. Only moves with at least 10 observations were considered, and four moves not in line with the predominant current were also excluded from the table.

| Current | Direction | Moves between stations | n | % of arrivals at the last station |  |  |  |
| --- | --- | --- | --- | --- | --- | --- | --- |
|  |  |  |  | 0:00-6:00 | 6:00-12:00 | 12:00-18:00 | 18:00-24:00 |
| With | NW > SE | P1,OW10 | 86 | 4.7 | 55.8 | 26.7 | 12.8 |
|  | NW > SE | OW10,P3 | 36 | 8.3 | 41.7 | 44.4 | 5.6 |
|  | NW > SE | OW3,OW5 | 15 | 6.7 | 40 | 26.7 | 26.7 |
|  | NW > SE | P3,P5 | 11 | 0 | 18.2 | 45.5 | 36.4 |
|  | NW > SE | OW3,P1 | 94 | 31.9 | 38.3 | 16 | 13.8 |
| Against | SE > NW | OW10,P1 | 239 | 41.8 | 3.4 | 17.6 | 37.2 |
|  | SE > NW | P1,OW3 | 61 | 55.7 | 11.5 | 11.5 | 21.3 |
|  | SE > NW | P3,OW10 | 39 | 28.2 | 2.6 | 17.9 | 51.3 |
|  | SE > NW | P3,OW10,P1 | 39 | 53.8 | 2.6 | 7.7 | 35.9 |
|  | SE > NW | P3,P1 | 26 | 69.2 | 7.7 | 3.6 | 19.2 |
|  | SE > NW | OW5,OW3 | 17 | 47.1 |  | 17.7 | 35.3 |

### Currents

In the Gulf area, the currents in the upper 40 m in the HYCOM model varied widely (Fig. 6). This general circulation pattern was consistent with the long-term currents in this region (Thoppil and Hogan 2012). However, within the acoustic array, the current direction was consistent during the whale shark season May–October, flowing to the southeast (Fig. 6).

**Fig. 6.**
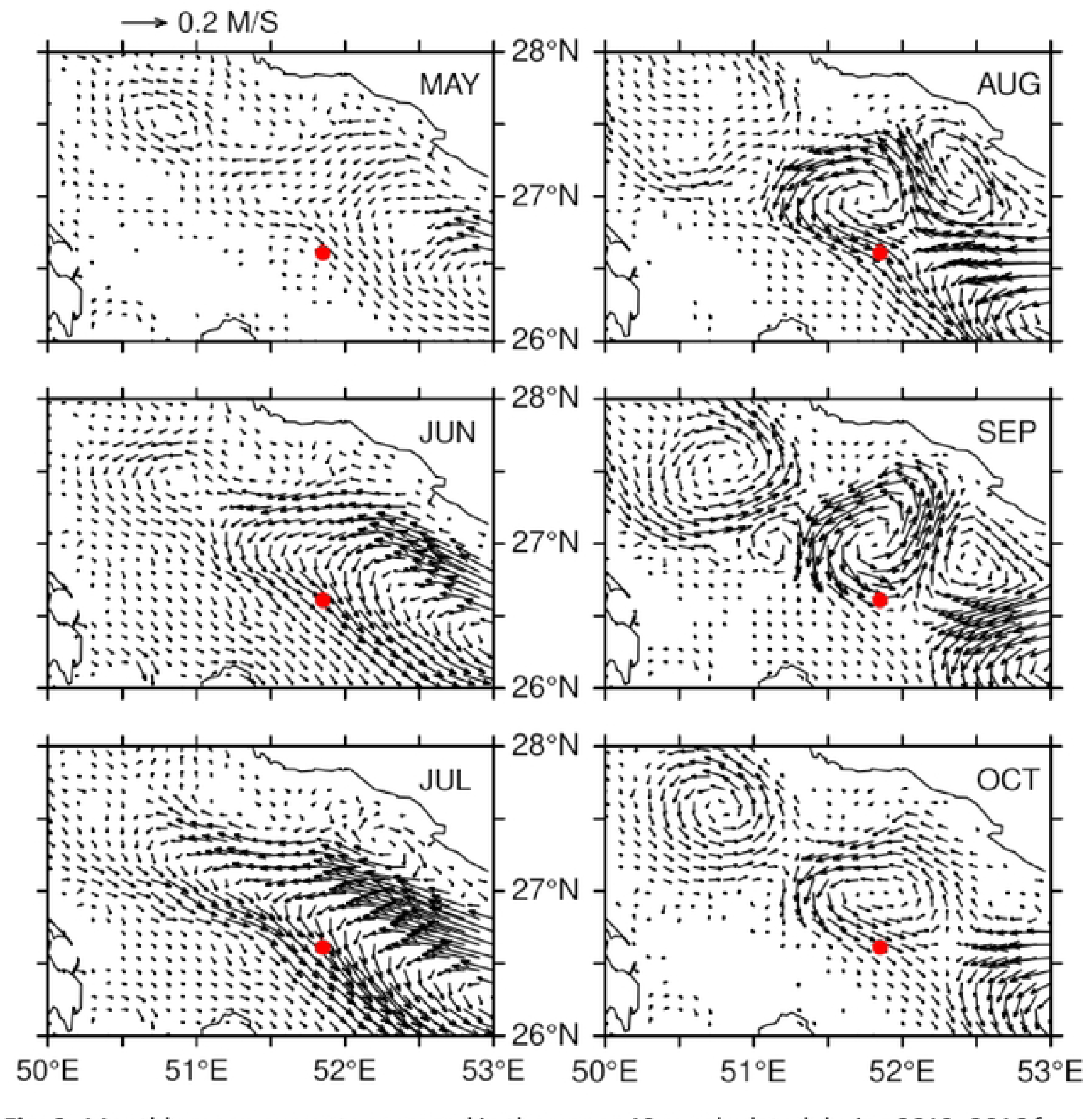
Monthly mean currents averaged in the upper 40 m calculated during 2012-2016 from a 1-km regional Hybrid Coordinate Ocean Model [HYCOM; 38]. The red circle indicates the location of the acoustic array.

## Discussion

Whale sharks exhibited a high degree of seasonal residency to a small core habitat area, adjacent to one of the gas platforms within the Al Shaheen field. Rather than being platform-associated, however, the sharks appear to be targeting a specific spawning area for mackerel tuna, with the tuna eggs providing a reliable and energy-dense food source. Individual whale sharks will repeatedly return to this spawning area each afternoon or overnight, then feed on drifting eggs between ∼7 am and ∼12 pm. Even though mean tag retention was 161 days, 27% of sharks were seen in two or more seasons, suggesting that a high proportion of whale sharks return each year; this is supported by the 41% inter-annual sighting rate of photo-identified sharks [36] and 58% of satellite-tagged sharks [31] during fieldwork at Al Shaheen from 2011–14. This area is clearly an important seasonal habitat for this globally Endangered species, emphasising the importance of spatial management for both whale sharks and the mackerel tuna spawning area.

### Residency

Whale sharks had high residency, with a combined R_max_ of 0.31 and mean of 0.56 ± 0.31. This means that whale sharks were detected on 31% of possible days. While R_max_ can overestimate residency for sharks that were only detected within a single season, it was still high at 0.21 for the 32 sharks that were detected in multiple seasons. The aggregation site off Qatar is thus clearly an important habitat for whale sharks in the region. Elsewhere, acoustic telemetry estimated a shorter combined R_max_ of 0.11 off St Helena [24] and 0.16 off Ningaloo Reef [21]. The constellations in Qatar and St Helena both comprise a high proportion of large, mature sharks, but St Helena appears to be less frequented, perhaps because feeding opportunities are rare there [24]. Whale sharks in the Red Sea constellation were often feeding and had a similar R_max_ to those in Qatar [0.26; 20], which suggests that residency is higher when feeding is the main driver of the aggregation. This is further supported by the passive acoustic study from Mafia Island in Tanzania, where whale sharks feed in high-density plankton patches, which found an even higher R_max_ of 0.39 [17,23].

Whale sharks off Isla Contoy in Mexico aggregate in large numbers to surface-feed on little tunny (*Euthynnus alletteratus*) eggs [41] in a similar manner to sharks at Al Shaheen, with tuna egg biomass at Al Shaheen sometimes even higher than the former [26]. The potential daily calorific intake of sharks at Isla Contoy is estimated to greatly exceed that needed for basal metabolic rate [18,42], or indeed the mean ration for whale sharks in aquaria [43], indicating that the tuna spawning event in Al Shaheen is a rich feeding opportunity for months at a time. This is reflected by the high residency we observed, with half of all sharks having consecutive daily detections for at least two weeks, and 12 individuals (10.3%) for over a month. However, there were also days during the main season with few whale sharks detected, when most were outside of the array. Some of them may have moved ∼120 km to a nearby location in Saudi Arabia where a secondary whale shark hotspot was identified in a satellite tagging study [31]. To better understand why whale sharks leave the area during the season, it will be important to assess fish egg density in the area over time as it is possible that tuna spawning fluctuates over the season.

### Sexual and size-based segregation

Like many whale shark aggregations [14], Al Shaheen is biased (∼69%) toward male sharks [36,44]. We tried to tag a more equal mix of male to female sharks in this study – 57% of the known-sex sharks were males – to facilitate a comparison of behaviour between the sexes. Females (combined R_max_ = 0.31) had a similar residency level to males (0.27). Although male sharks have a larger mean TL (7.25 m) than females (6.44 m) at Al Shaheen [36], this did not appear to influence the results observed in the present study. One of the hypotheses put forward for sexual segregation in whale sharks is a desire for females to avoid male harassment, in the form of undesired courtship or mating attempts [14]. Our residency data suggest that either this is not an important driver, or alternatively that the benefits of the rich food source outweigh any drawbacks for the female sharks that do choose to frequent the site.

### Habitat use

Satellite tracking showed that the Al Shaheen area is the hotspot of whale shark activity within the Gulf [31]. Here, we expand on this by identifying the small core habitat within this area, located around station P1. This station recorded most detections (65%) and almost all tagged sharks (97%). The location of the activity hotspot was stable, both within the season and across years. As whale sharks aggregate at Al Shaheen to feed on fish eggs [26], this indicates that the mackerel tuna consistently spawn in the same area. While the exact location and extent of the mackerel tuna spawning ground is yet to be established, our results indicate that it is near platform P1, where most whale sharks were detected, or slightly north-west (upcurrent) of it. There are distinct hydrological characteristics in the area that could explain the general area of the spawning ground. Hydrographic fronts in the area are formed by mesoscale cyclonic eddies carrying Indian Ocean water along the Iranian coast [38]. At the same time, water from the northern part of the Gulf is transported along the Saudi Arabian coast to the northern tip of Qatar [38]. It is possible that these currents are mixing in the Al Shaheen area, resulting in increased productivity and food availability for the spawning mackerel tuna. Finding the spawning ground and ensuring protection from potential fisheries exploitation will be important to help the tuna stock in the Gulf and protect this important feeding area for whale sharks. Even though the exact spawning area has not been identified at this stage, the small extent of this area means that spatial management could proceed with the information available now.

### Association with platforms

Whale sharks were detected on receivers at oil and gas platforms and on receivers in open water. Station P1, located at a two-platform structure connected by a gangway, was the centre of the relatively small whale shark core habitat and had by far the most detections. Open water station OW10 ∼4 km southeast of P1 registered the second-most detections and was on the edge of the core habitat. OW stations 3, 5, and 7 were also located within 4 km of P1 but recorded fewer detections than P1 and OW10. The only platform with significant numbers of detections other than P1 was P3, located southeast of P1 and OW10. Considering that many of the receivers at other platforms had few detections, and that the distance from the centre of the aggregation was the primary explanatory variable in the GLM, our results suggest that the whale shark aggregation is driven by the feeding opportunity rather than the presence of platforms. In the wider Gulf area, there are more than 800 platforms [29] and we are not aware of any others at which whale sharks are regularly seen, let alone to aggregate near as they do around P1 in Al Shaheen. A second likely whale shark aggregation in the Gulf is ∼120 km to the north-west of the Al Shaheen oil field, close to the reef plateau of the Rennie Schoals in Saudi Arabia waters [31]. Here, the nearest platform is at the Abu Sa’afa field, ∼10 km distant.

It remains unclear whether the spawning tuna aggregate here only since the platforms were constructed or have done so before artificial structures were built at Al Shaheen. Other studies suggested that it is unlikely for tuna to alter their spawning locations in response to structures being built, but this was based partly on the assumption of their short-term association with structures [2]. By contrast, tuna spawn for several months at Al Shaheen. The relatively shallow bathymetry of the Gulf may not provide other typical aggregation areas for tuna, such as seamounts [45], which could further increase the importance of these human-made structures for tuna spawning here. A more detailed analysis of the ecology and spawning behaviour of mackerel tuna at Al Shaheen is needed to better understand their relationship with structures and their influence on whale sharks.

### Seasonality

The results from this passive acoustic telemetry study confirm the seasonality observed for this aggregation in visual surveys and with satellite tags [26,31]. The main whale shark season at Al Shaheen is from May–September, with additional high activity in April and October in some individual years. During November–March, few whale sharks were detected. As such, passive acoustic telemetry confirmed the pattern seen with other methods and this aggregation is distinctly seasonal. A similar result was seen for whale sharks at Shib Habil in the Red Sea [20], while passive acoustic telemetry revealed cryptic residency in whale sharks throughout the year at Mafia Island in Tanzania [19] and their year-round presence at Ningaloo Reef in Australia [21]. It is plausible that, in general, whale shark aggregations driven by fish spawning are seasonal, as fish spawning is also seasonal, for example during Apr–Sep for mackerel tuna in southwestern India [46], particularly if no other whale shark prey are available near these locations during other times of the year. The start and duration of fish spawning may be influenced by biophysical variables, leading to years with high detections earlier or later than the main whale shark season, and a better understanding of the drivers of mackerel tuna spawning at Al Shaheen will likely allow predicting when whale sharks can be seen feeding at this site.

### Temporal trends

Visual observations showed that whale sharks are feeding at the surface from ∼7am to ∼12pm, suggesting that they may leave the Al Shaheen area until the next morning [26]. Satellite tags also showed that they stay shallower in the morning and swim deeper during the rest of the day and night when they are at Al Shaheen, indicating that they only surface feed during the morning [31]. Here, we found no difference in detections between the day and night, clarifying that the sharks stay within the area covered by the acoustic stations.

We also found that the peak in relative detections moved with the predominant current over the course of the day, starting in the north-west (stations OW3, P1) during the night and early morning and ending in the south-east in the afternoon (P3, P5), before the sharks swam against the current. Detections increased again at OW3 and P1 from late afternoon and over the night, indicating that whale sharks return to the location in anticipation of the spawning. Movements, defined as being detected at a subsequent station within 3h of first being detected at a station, were largely with the current during the day between 6am–6pm and against the current during the night. The HYCOM current model showed that the flow was south-easterly throughout the whale shark season months in the years of our study 2012–2016.

Due to the high water temperatures at Al Shaheen (26–35 °C) during the whale shark season, the eggs likely develop and hatch quickly. Using the equation in Pauly and Pullin [47] and an egg diameter of 0.88 mm (D. Robinson unpubl. data), mackerel tuna eggs hatch within 0.55–0.89 days (13.2–21.4 h) during the whale shark season. Combined, it appears that mackerel tuna spawn in the late hours of the night, and that the whale sharks feed on fish eggs which floated to the surface in the early morning, drift with the floating eggs at the surface towards the south-east during the morning and early afternoon, and then cool off at depth during the night and return to the north-west anticipating the next spawning event.

### Notes on management

Whale sharks are Endangered, having suffered severe population decline over the past three generations. They face a variety of potential threats in the Gulf region [48], likely making this one of the more at-risk populations for the species [49]. Off Oman, whale sharks are still taken by fishers (D. Robinson unpubl. data), and the whole area – particularly the Strait of Hormuz – has intense shipping traffic which could severely impact the sharks in this region [50]. Surface-feeding whale sharks within the Al Shaheen area are at high risk from boat strike, so the specific movement data presented here provide an opportunity for careful spatial management of vessel movements in the Al Shaheen field through the whale shark season.

## Acknowledgements

We thank everyone involved in the Qatar Whale Shark Research Project, as well as the staff at the Qatar Ministry of Municipality and Environment (QMME) and the Qatar Coast Guard for providing the platform to carry out field research in Qatar. We thank Maersk Oil (A.P Møller Mærsk) for providing the majority of financial support for the purchasing of equipment and the staff of the Maersk Oil Research and Technology Centre. We thank Sheikh Abdulah bin Nasser bin Khalifa Al Thani for supporting the Qatar Whale Shark Research. We also thank the offshore platform workers for their dedicated support with data collection.

## Notes

### Competing Interest Statement

The authors have declared no competing interest.

